# Single molecule imaging of the central dogma reveals myosin-2A gene expression is regulated by contextual translational buffering

**DOI:** 10.1101/2024.02.11.579797

**Authors:** O’Neil Wiggan, Timothy J. Stasevich

## Abstract

While protein homeostasis is a hallmark of gene regulation, unraveling the hidden regulatory mechanisms that maintain homeostasis is difficult using traditional methods. To confront this problem, we CRISPR engineered a human cell line with multiple tags in the endogenous MYH9 gene, which encodes the essential and ubiquitous myosin-2A cytoskeletal motor. Using these cells, we imaged MYH9 transcription, translation, and mature mRNA and protein in distinct colors, enabling a full dissection of the central dogma. Our data show that MYH9 transcription is upregulated in an SRF-dependent manner in response to cytoskeletal cues and that MYH9 translation can either buffer or match the transcriptional response depending on context. Upon knockdown of actin-depolymerizing proteins like cofilin, translation efficiency drops by a factor of two to buffer strong transcriptional upregulation, likely to help prevent excessive myosin activity. In contrast, following serum stimulation, translation matches the transcriptional response to readily reestablish steady state. Our results identify contextual translational buffering as an important regulatory mechanism driving stable MYH9 expression. They also demonstrate the power and broad applicability of our cell line, which can now be used to accurately quantify central dogma dynamics in response to diverse forms of cellular perturbations.

## Introduction

A grand challenge in cell biology is to image the full central dogma as it plays out in living cells, one molecule at a time. Although tags now exist to separately image individual proteins, mRNA, and sites of transcription and translation ^1–4^, the tags are rarely combined into a single experimental system and even more rarely incorporated into endogenous genes ^5^. This has made it difficult to pinpoint at what level gene regulation occurs ^6^; for example, does a decrease in total protein reflect less translation or more degradation? Alternatively, if levels remain unchanged, does that mean that transcription and translation are also unchanged, or could they instead be changing together, but in opposition?

The MYH9 gene exemplifies this challenge. MYH9 encodes non-muscle myosin-2A heavy chain, a motor protein that assembles into bipolar filaments that use ATP to power contraction of the actomyosin cytoskeleton ^7^. This contraction generates intracellular forces that govern cell architecture, migration, adhesion, and division. MYH9 is essential and, like beta-actin, is sometimes categorized as a housekeeping gene, implying stable expression and minimal regulation ^7, 8^. Consistent with this, myosin activity is thought to be mainly regulated by post-translational phosphorylation, a dynamic and reversible mark that activates myosin motors ^9^. On the other hand, the MYH9 gene has also been categorized as both an oncogene and tumor suppressor ^10, 11^, and some studies have even shown cancer metastasis relies on MYH9 upregulation ^12^. These studies highlight the need to better understand MYH9 regulation and they suggest both too little and too much MYH9 expression is detrimental. This raises an important and generalizable question: do multiple competing regulatory mechanisms strictly enforce the stable expression of genes like MYH9, or does the stability instead arise from a lack of regulation?

To directly address this question and better understand how gene expression homeostasis can be established and maintained, we engineered a unique human cell line with multiple tags knocked into the endogenous MYH9 gene. By co-tracking MYH9 transcription, translation, and total mRNA at the level of single molecules, we show that translational regulation is a critical aspect of MYH9 expression, one that can be independently regulated from transcription. During cell division or upon the loss of actin depolymerizing proteins, translation buffers transcriptional upregulation, presumably to offset detrimental myosin-2A overexpression. In contrast, upon a release from serum starvation, translation increases in sync with transcription to quickly establish cellular phenotypic changes. Collectively, our results highlight the complexity of the central dogma and provide compelling evidence that even apparently stable genes like MYH9 can hide multitiered and context-dependent gene regulatory mechanisms.

## Results

### A system to co-image endogenous transcription and translation with single-molecule precision

To evaluate gene expression for endogenous MYH9 in human HeLa cells, we performed sequential rounds of CRISPR Cas9 gene editing. First, we incorporated an in-frame C-terminal mClover tag (a GFP variant referred to herein as GFP) at the end of exon 40 followed by 24×MS2 stem loops upstream of the native 3’ untranslated region. Next, we integrated an N-terminal 6×Flag tagged mCherry after the start codon (**Figs. 1A, S1A**). Following successive rounds of gene editing, we isolated subclones where both MYH9 alleles were C-terminally tagged and one was N-terminally tagged (**Fig. 1B**). The tags did not appear to impair MYH9 function because Myo2A-GFP localized to actomyosin structures, including cortical bundles and stress fibers (**Fig. 1C**) and MYH9 mRNA levels were not significantly different than unedited cells (**Fig. S1B**). Using these cells we could visualize mature Myo2A proteins in green (GFP), nascent Myo2A peptide chains at translation sites in red (using Cy3-conjugated anti-Flag antibodies), and Myo2A-endoding mRNA in far-red (using Halo-tagged MS2 coat protein with JF646 ligand), enabling a full dissection of the central dogma in separate colors. Hereafter we refer to this cell line as eMyo2AGFP.

**Fig. 1.**
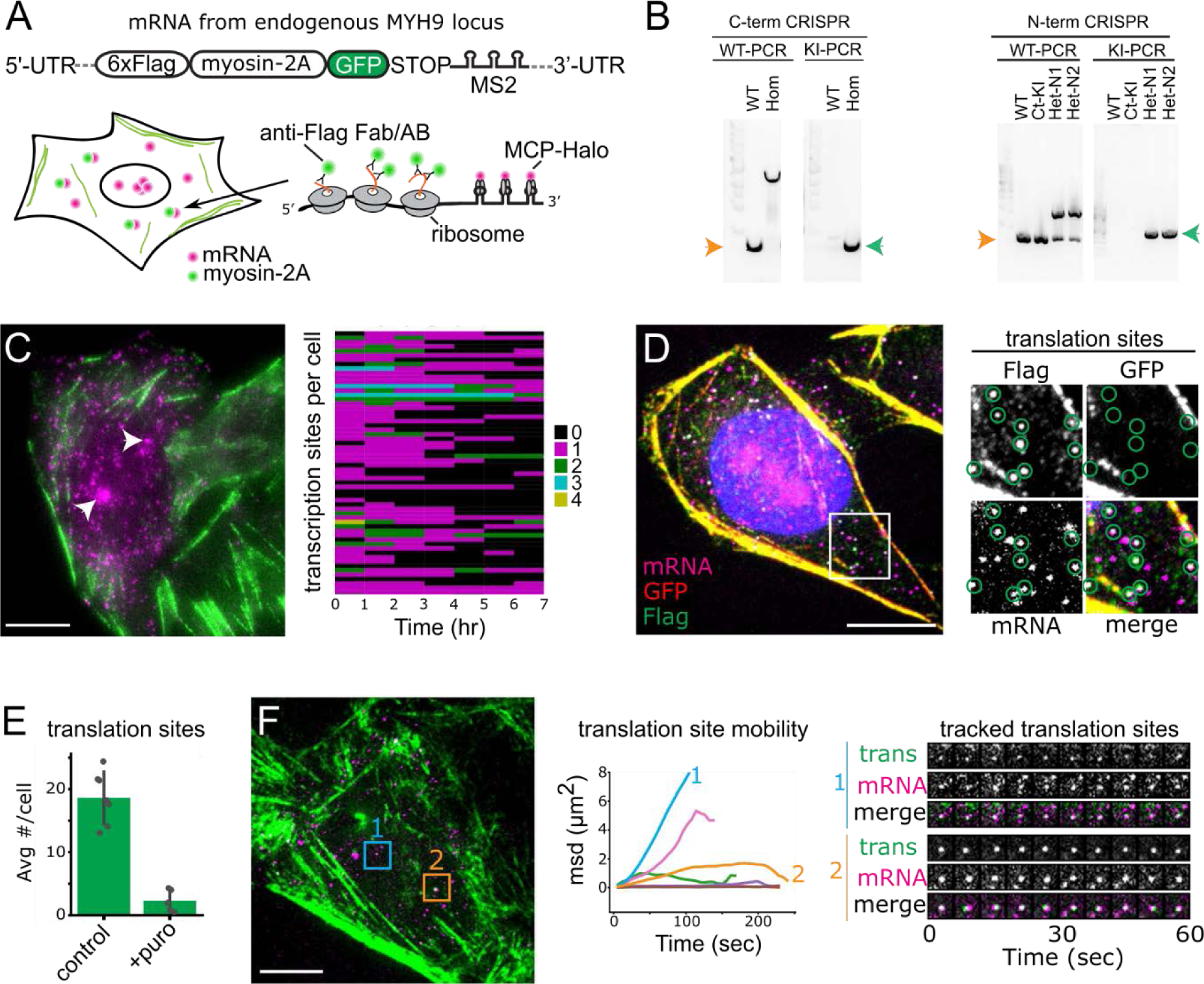
A cell line for dissecting the central dogma with single-molecule precision. **A.** Top, schematic of a tagged MYH9 transcript in CRISPR modified eMyo2AGFP cells. Below, cartoon of an eMyo2AGFP cell expressing myosin2A-encoding mRNA (magenta), myosin2A protein (green) and translation sites (colocalized foci of mRNA and protein). **B.** Genomic PCR for detection of wild type (orange arrow) or knock-in (green arrow, homo or heterozygous) alleles, following successive rounds of CRISPR editing. **C.** Live cell image of eMyo2AGFP cell with myo2A-GFP protein (green) and mRNA (magenta), arrowheads show mRNA transcription foci. Heatmap of numbers of transcription sites over 1hr intervals, each row corresponds to one cell (right). **D.** Representative confocal fixed cell image showing myo2A mRNAs without translation signals (magenta), and translating mRNAs marked by colocalized mRNA and Flag-myo2A (green) but absence of mature myo2A-GFP (red). Boxed region is enlarged (right) with translation sites encircled. **E.** Quantification of loss of translation sites following puromycin treatment. Mean ± SD, 580 cells control and 285 cells puro. **F.** Live cell confocal image of anti-FLAG Fab labeled Flag-myo2A (green) and mRNA (magenta). Mean squared displacement (msd) plots for 7 mRNA tracks (center), including translation site mobility of a single mRNA (1) or an mRNA cluster (2) for boxed regions also shown in time series images, right. Scale bars 10 μm.

To demonstrate our ability to detect both transcription and translation with single molecule sensitivity in eMyo2AGFP cells, we first focused on steady-state MYH9 expression. Live imaging revealed bursts of transcription at between one and four sites per cell (**Fig. 1C and Movie S1**), as would be expected for a diploid locus. Cells had tens to hundreds of MYH9 transcripts in the nucleus and cytoplasm (**Fig. 1C**). To identify which were translated, we immunostained fixed cells and looked for bright FLAG signals that overlapped with mRNA yet lacked GFP (since GFP takes time to mature after translation). According to this metric, many cytoplasmic mRNA were translation sites (**Fig. 1D**) and these reassuringly disappeared upon treatment with the translational inhibitor puromycin (**Fig. 1E**).

To further characterize individual Myo2A translation sites, we performed live imaging with anti-FLAG intrabodies ^4^. This revealed diverse behaviors, from fast unidirectional movement to slow, nearly immobile diffusion (**Fig. 1F and Movie S2**). We usually detected translation from single mRNAs, although a small fraction (<1%) were in bright clusters, reminiscent of translation factories (**Fig. 1F, spot 2**) (ref 13). Altogether, our data demonstrate eMyo2AGFP cells are versatile tools to comprehensively examine broad aspects of MYH9 gene expression with single molecule precision.

### *M*YH9 translation efficiency is high and undergoes regulation during cell division

Confident in our ability to detect individual mRNA and translation sites, we next set out to quantify their average numbers in cells. We used fixed cells for this analysis because we obtained better signal to noise and tag saturation than is possible in live cells. On average, eMyo2AGFP cells contained 90 ± 47 total mRNA per cell (**Fig. 2A**). Of these, 53 ± 29 were in the cytoplasm, 20 ± 10 of which were being translated (we did not detect nuclear translation). Separately, the fraction of cytoplasmic mRNAs translating the FLAG tag was measured at 46 ± 6% (**Fig. 2B**). Since we only expect half of the cytoplasmic transcripts to encode the FLAG tag in the first place (due to its heterozygous insertion), we estimate the translation efficiency to be double our measured cytoplasmic fraction ∼ 93% (**Fig. 2B, projected**). These data therefore suggest that the majority of cytoplasmic MYH9 transcripts were being translated.

**Fig. 2.**
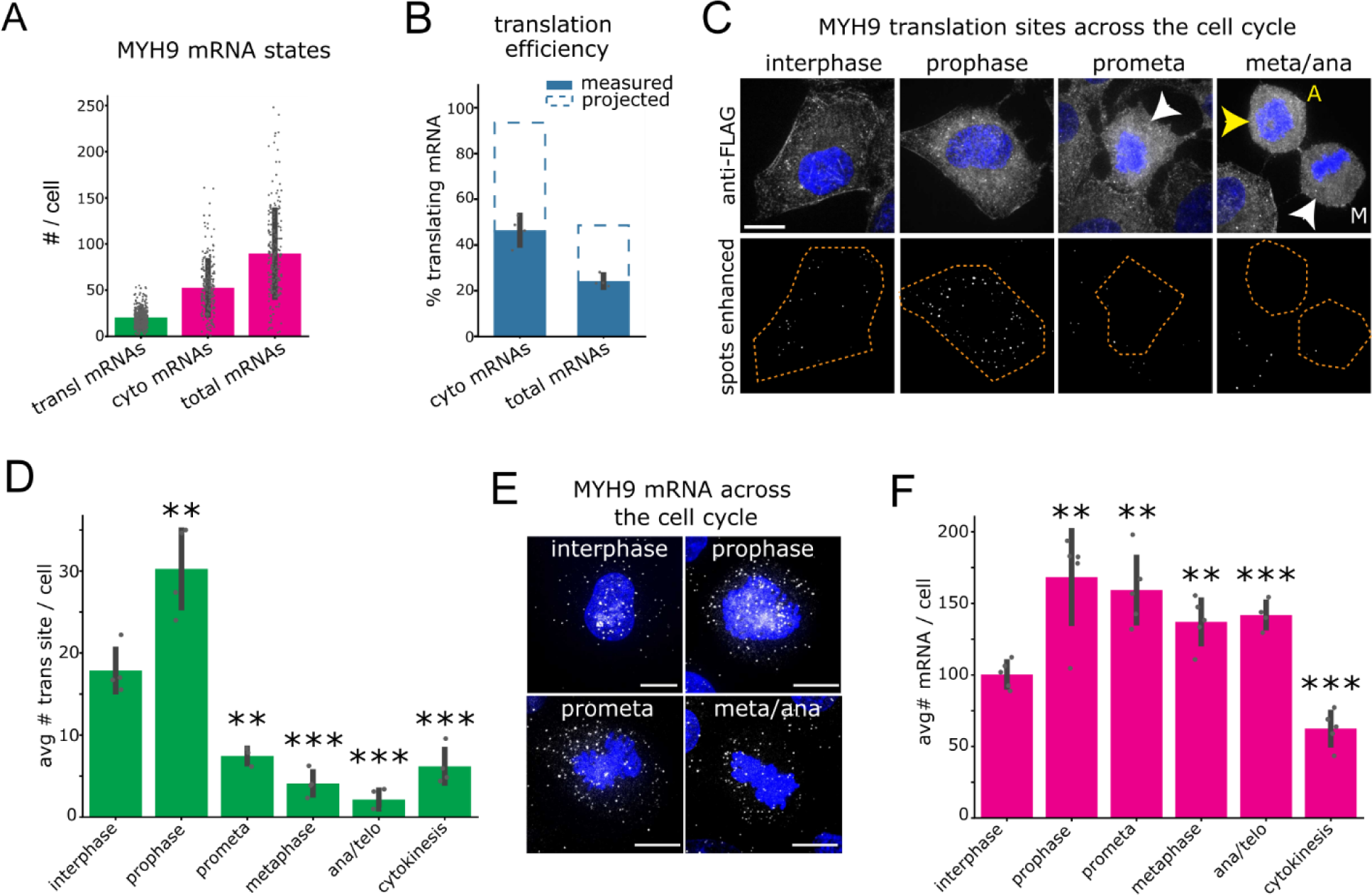
MYH9 translation is downregulated during cell division. **A**. Quantification of mRNAs and translation. Mean ± SD. **B.** Quantification of translating mRNA percentage. Note, in eMyo2AGFP cells there are two labeled MYH9 mRNA alleles, of which only one detects translation (measured). Mean ± SD, n= 361 cells. Dashed line (projected) shows projected % translation accounting for two alleles. **C.** Representative confocal immunofluorescence images of MYH9 translation sites labeled by anti-Flag and for DNA (blue). A-anaphase cell, M-metaphase cell. Lower panels show spot fluorescence enhanced by a LOG filter for outlined cells. **D.** Quantification of translation sites across the cell cycle. Mean ± SD, interphase n= 312 cells, mitotic n ≥ 20 cells/phase. **E.** Fluorescence images of MYH9 mRNAs labeled by Halo-MCP were imaged in fixed cells at different stages of the cell cycle (DNA in blue). **F.** Quantification of the average number of mRNA per cell. Mean ± SD, interphase n= 122 cells, mitotic n ≥ 18 cells/phase. Scale bars 10 μm. **p ≤ 0.01, ***p ≤ 0.001, Welch’s t-test.

The high efficiency of MYH9 translation is perhaps not surprising given the abundance of myosin-2A required to form the actomyosin cytoskeleton. However, this left us wondering if MYH9 translation is subject to regulation. To test this, we imaged eMyo2AGFP cells across the cell cycle (**Fig. 2C**), avoiding the use of drugs commonly used to synchronize cells (which can suffer from artifacts ^14^). We hypothesized translation might be modulated during cell division since myosin-2 motors are known to play an important and highly conserved role in contractile ring assembly. Our analysis confirmed this: we observed a sharp increase in the number of translation sites at prophase relative to interphase, followed by a dramatic drop from prometaphase through cytokinesis (**Fig. 2D**). Interestingly, while the initial increase could be explained by a commensurate increase in MYH9 mRNA levels, the later drop did not follow the mRNA pattern (**Fig. 2E, F**). Thus, it appears that MYH9 translation is downregulated independently from transcription just after prophase.

Rounded mitotic cells generally have reduced cell surface area relative to those of interphase (**Fig. S2A**) and there was a strong positive correlation between cell size and MYH9 translation (**Fig. S2B**), so we explored whether cell size alterations could account for the reduced translation. However, normalization of MYH9 translation sites to cell area still showed reduced MYH9 translation for cells in prometaphase onwards (**Fig. S2C**). We conclude that reduced mitotic MYH9 translation is not a function of decreased cell areas. In summary, our results identify existence of a post-transcriptional mechanism that modulates the overall high efficiency of MYH9 mRNA translation in a cell-cycle dependent manner.

### Silencing of actin depolymerizing proteins stimulates SRF-dependent MYH9 transcriptional bursts

Having characterized MYH9 expression in steady state, we now turned our attention to MYH9 stimulation. In an earlier study we showed that knockdown of two closely related actin depolymerizing proteins, cofilin and ADF, results in detrimental myosin contractile activity ^15^. Over a period of 72 hours, this leads to excessive forces in cells that contort nuclei into tortured shapes that are reminiscent of those founds in tumors (**Fig. 3A**). This detrimental phenotype can be rescued by co-inhibiting myosin during cofilin/ADF silencing, demonstrating the phenotype is due to excessive myosin-based forces (**Fig. 3B, and ref 16** for quantification). We previously showed this is in part due to enhanced association of myosin with F-actin (since cofilin competes with myosin for F-actin). However, *in vivo* studies of ADF null mice suggested a link between defective actin dynamics and actomyosin gene expression ^17^. This led us to ask if deregulated MYH9 expression was also partly responsible.

**Fig. 3.**
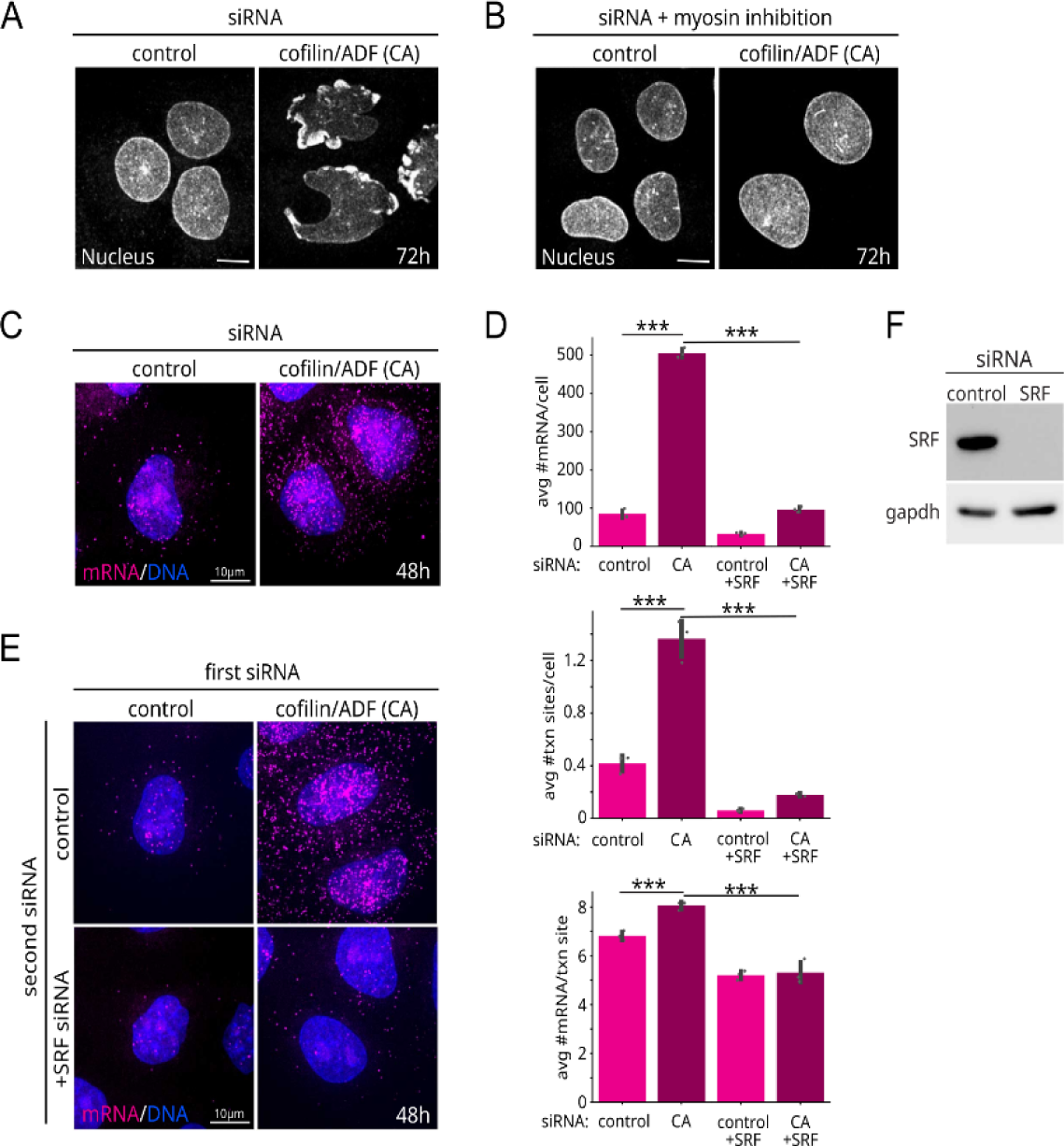
SRF-dependent MYH9 transcriptional bursts in response to knock-down of actin polymerizing proteins. **A, B**. Representative immunofluorescence images of nuclear envelope labeled cells depicting nuclear dysmorphology induced by cofilin/ADF silencing (A) and rescue by myosin inhibition via Y27632 treatment (B). **C-E**. MCP-Halo fluorescence images of MYH9 mRNA (magenta, C and E) and quantification of MYH9 mRNAs (D, top), transcription sites (D, middle) and burst amplitude (D, bottom) following cofilin/ADF silencing (C) and co-silencing of SRF (E). Scale bars 10 μm. Values are mean ± SD, n ≥ 186 cells/treatment. *** p ≤ 0.001, Welch’s t-test. **F**. Immunoblot of SRF siRNA silencing.

To test this, we silenced cofilin/ADF expression using siRNA in eMyo2AGFP cells (**Fig. 3C**). Consistent with MYH9 deregulation, the number of MYH9 mRNA per cell dramatically increased from ∼100 to ∼500 after 48 hours (**Fig. 3D, top, Control & CA**; note at 72 h, when nuclei are contorted, there were too many mRNA to count). The mRNA increase was caused by a roughly 3-fold increase in the number of transcription sites per cell (burst frequency; **Fig. 3D, middle, Control & CA**) in combination with a modest increase of ∼1.3 mRNA per transcription site (burst amplitude; **Fig. 3D, bottom, Control vs CA**). Live-cell analysis confirmed increased temporal MYH9 burst activity for cofilin/ADF silenced cells (**Fig. S3C**). To see if there were additional changes to transcription kinetics after an allele is activated, we tracked active sites over a period of 2-5 hours (**Fig. S3A and Movie S3**). However, classification of their fluctuations using a three-state Hidden Markov Model ^18, 19^ revealed no significant changes (**Fig. S3B**). It remains possible that measurements over periods longer than made for our study, would reveal changes to heterogenous long refractory transcriptional OFF times, as suggested by a recent study ^20^. A likelihood, given we detected some cells with extended OFF intervals (**Fig. S3C**). Together, these data suggest the major impact of cofilin/ADF knockdown on MYH9 transcription is enhanced activation.

Stimulation of transcription via simultaneous frequency and amplitude modulation is not universal ^21^, but has been observed for genes such as beta-actin ^22^, whose transcription is linked to a master cytoskeleton transcription factor, Serum Response Factor (SRF) ^23^. As MYH9 is also a putative target of SRF, we wondered if SRF is responsible for the enhanced transcription. To see if this was the case, we repeated our cofilin/ADF knockdown experiments, but in addition we co-silenced SRF (**Fig. 3E, F**). Remarkably, this almost completely blocked the increase in MYH9 mRNA levels, attenuating both burst frequency and amplitude (**Fig. 3D, Control+SRF & CA+SRF**). These results identify SRF as an essential mediator of transcription burst dynamics for the MYH9 gene in both basal and stimulated conditions. The data also argue that elevated MYH9 mRNA levels following cofilin/ADF loss results from increased transcription and not to other possibilities, such as mRNA stabilization.

### MYH9 transcription is linked to cytoskeletal signaling that promote MKL1 nuclear localization

By what mechanism does loss of cofilin/ADF lead to SRF-dependent MYH9 transcription? This is an important question because cofilin inactivation, just like myosin-2A overexpression, has been linked to cancer ^24^. Transcription of cytoskeletal genes by SRF relies on MKL-family cofactors (a.k.a. MAL, MRTF, ref 23). According to a model (**Fig. 4A**), when G-actin subunits assemble into actin filaments (F-actin), MKL can enter the nucleus with SRF and activate cytoskeletal genes. However, if G-actin levels become sufficiently high, they bind MKL and prevent its translocation into the nucleus, causing attenuation of cytoskeletal gene expression. This cascade ostensibly creates a self-regulatory feedback loop, dictated by the balance of G-to F-actin.

**Fig. 4.**
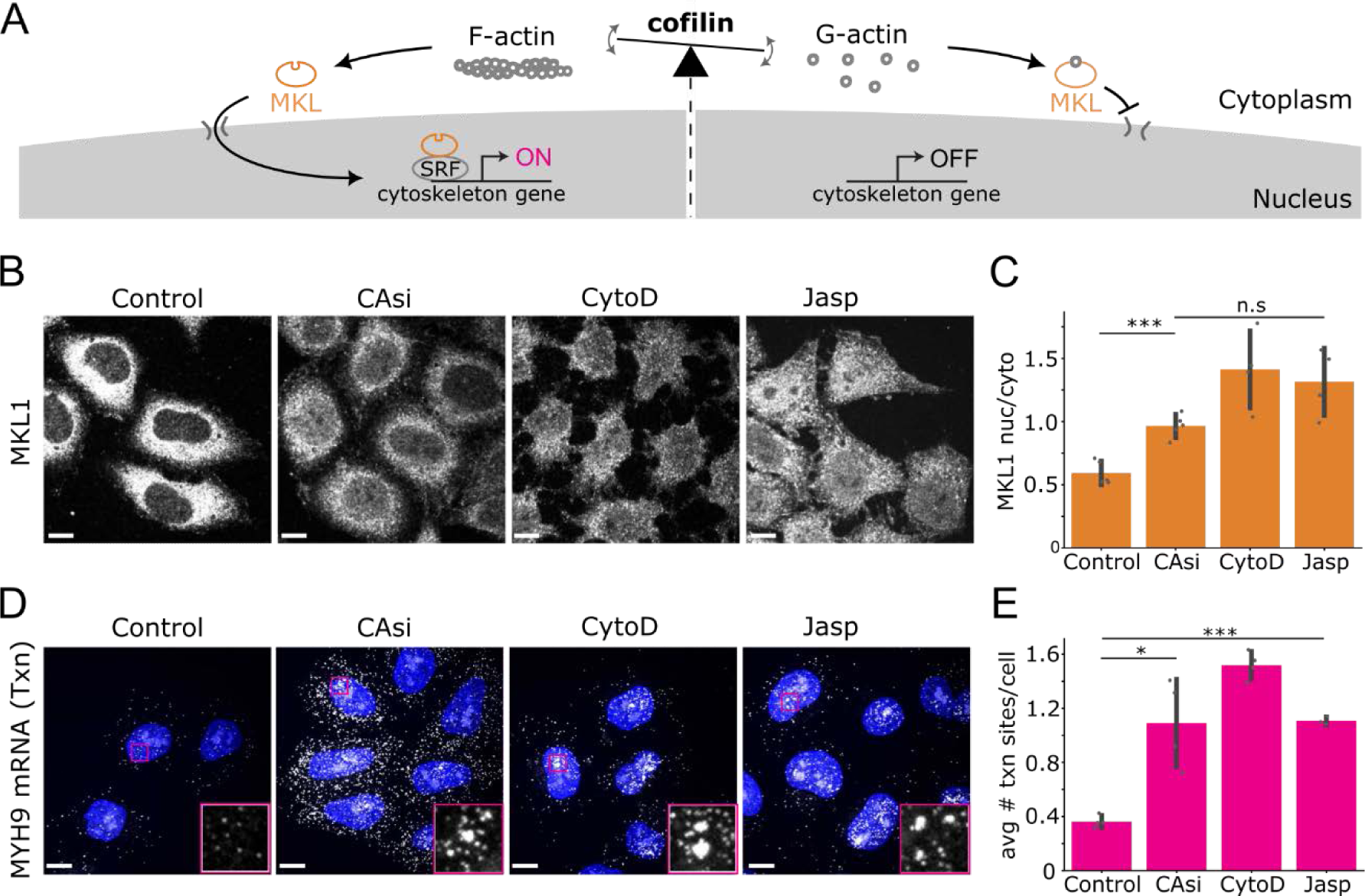
Altered cytoskeletal dynamics promote MKL1 nuclear localization and MYH9 transcriptional bursts. **A**. Illustration of model by which cofilin activity and actin dynamics may regulate MKL localization and SRF-dependent cytoskeletal gene expression. **B and C**. Nuclear level confocal immunofluorescence images (B) and quantification (C) of MKL1 nuclear/cytoplasmic distribution for cells treated with cofilin/ADF siRNAs (CAsi), cytochalasin D (cytoD) and Jasplakinolide (Jasp). **D and E**. Confocal fluorescence images (D) and quantification (E) of MCP-Halo labeled MYH9 mRNAs. DNA in blue. Insets (D) show enlargement of boxed regions depicting transcription sites. Note, treatment periods are 48h for siRNAs and 2h for drugs. Scale bars 10 μm. Values are mean ± SD, n ≥ 350 cells/treatment (C), n ≥ 330 cells/treatment (E). *p < 0.05, ***p ≤ 0.001, n.s not significant by Welch’s t-test.

With this model in mind, since cofilin and ADF both disassemble actin filaments into G-actin, their loss should lower G-actin levels, causing MKL to translocate into the nucleus and activate cytoskeletal genes like MYH9. To test this model, we measured MKL1 levels in eMyo2AGFP cells at 48 hours post cofilin/ADF silencing and indeed found a significant increase in the nuclear fraction of MKL1 (**Fig. 4B, C**). However, the increase was not particularly striking, as the majority of MKL1 remained in the cytoplasm. We saw a similar result in a set of related experiments in which we promoted levels of MKL unbound by G-actin using drugs (cytochalasin D displaces G-actin from MKL and jasplakinolide stabilizes F-actin ^25^, reducing G-actin) (**Fig. 4B, C**). In all cases, MKL1 nuclear localization was only modest despite a robust SRF-dependent MYH9 transcriptional response (**Fig. 4D, E**). Our data therefore point to a feedback mechanism between actin cytoskeletal organization and MYH9 transcription and suggest that the threshold of nuclear MKL1 levels necessary for activation of endogenous SRF/MKL targets may be lower than commonly expected.

### MYH9 expression is stabilized by a translational buffering system

Given the fact that persistent MYH9 upregulation leads to excessive intracellular forces that contort cell nuclei and possibly contributes to cancer, we wondered if there were any post-transcriptional expression mechanisms that might counter the strong transcriptional induction seen after cofilin/ADF silencing. As a first check, we plotted total myosin-2A protein levels versus the number of cytoplasmic mRNA in eMyo2AGFP cells. This revealed a positive correlation in control conditions, but only a weak correlation after cofilin/ADF silencing (Pearson r = 0.61 vs 0.11, **Fig. 5A**). Regression slopes indicated a reduced rate of change of MYH9 protein to mRNA (slope = 2.3×10^7^ vs 2.9×10^6^), suggesting some form of protein buffering, either at the level of mRNA translation or protein decay.

**Fig. 5.**
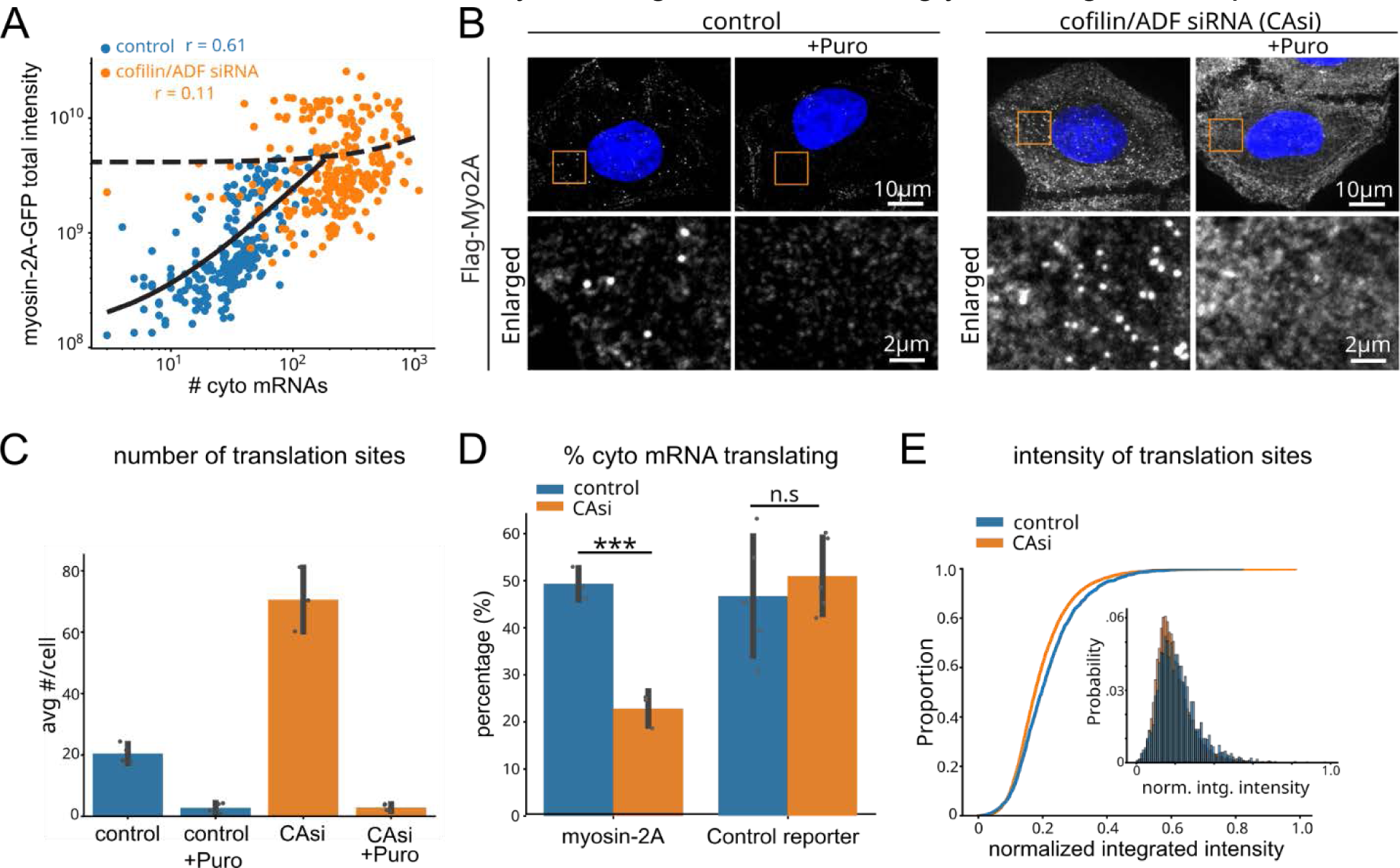
Depletion of actin depolymerizing proteins results in MYH9 translational buffering. **A.** Log scale correlation plot of per cell cytoplasmic MYH9 mRNAs vs total myosin-2A-GFP intensity with control or cofilin/ADF siRNA (CA) treatments. **B and C**. Confocal immunofluorescence images (B) and quantification (C) of anti-Flag labeled MYH9 translation sites. Boxed regions are shown in enlargements. Values are mean ± SD, n ≥ 201 cells/treatment. **D.** Proportion of translating myosin-2A or KDM5B control mRNAs. Mean ± SD, n ≥ 209 cells/treatment. **E.** Cumulative distribution functions and histogram (inset) of normalized translation site intensity, control n=165 cells and 2098 translation sites, cofilin/ADF siRNA (CA) n = 174 cells and 7272 translation sites. ***p ≤ 0.001, n.s, not significant, Welch’s t-test.

To see which of these was at play, we quantified the number and intensity of translation sites in control and cofilin/ADF silenced cells. Knockdown of cofilin/ADF resulted in a 3.5-fold increase in MYH9 translation sites (**Fig. 5B, C**). Consistent with translational buffering, the fraction of cytoplasmic mRNA that were being translated was reduced by a factor of more than two (**Fig. 5D**). We did not detect a similar decrease in translation efficiency for a control translation reporter, HA-KDM5B-MS2 (**Fig. 5D**), suggesting the reduction in translation efficiency is not global. Interestingly, although we expected the intensity of individual MYH9 translation sites to also be reduced, this was not the case (**Fig. 5E**). Instead, their intensities remained similar, suggesting whatever form MYH9 translational buffering takes, it operates more like an on/off switch than a dimmer. In conclusion, our data demonstrate MYH9 is regulated by a translational buffering system that stabilizes MYH9 expression under conditions of persistent transcriptional upregulation, presumably to delay the detrimental effects of myosin-2A overexpression.

### MYH9 transcription and translation cooperate upon serum stimulus and are both required for actomyosin remodeling

Given our previous results we wondered if translational buffering is a universal feature of MYH9 expression. Growth factor stimulation produces rapid (< 30 min) actomyosin cytoskeletal reorganization concurrent with SRF transcriptional activation ^26–28^. How transcription is coupled with translation to this physiologic cue, and whether new protein synthesis may contribute to the near-term cytoskeletal response remain unknown.

To address these questions, we serum starved eMyo2AGFP cells and then re-exposed them to serum, measuring total MYH9 mRNA and protein production (**Fig. 6A, B**). This led to rapid actomyosin rearrangements, including the formation of circumferential actomyosin bundles after one hour and an increase in the numbers of ventral stress fibers over the central portion of cells over a period of three hours (**Fig. 6A**). Serum stimulation produced a transient elevation of MYH9 transcription that peaked at 1h post stimulation, as evidenced by an increase in the burst fraction (**Fig. 6C, S4D**). Unlike the translational buffering we observed upon cofilin/ADF silencing, serum stimulation did not significantly alter the fraction of translating cytoplasmic mRNAs (**Fig. 6C**). Instead, the numbers of MYH9 translation sites continued to increase in tandem with the number of cytoplasmic mRNA (**Fig. 6C, S4G**). Furthermore, a modest positive linear correlation between MYH9 protein and mRNA levels was maintained at this post-stimulus timepoint (**Fig. S4A**, Peason r = 0.37 vs 0.43). Altogether, these data suggest that MYH9 translation does not always counter the transcriptional response, implying translational buffering is context-dependent.

**Fig. 6.**
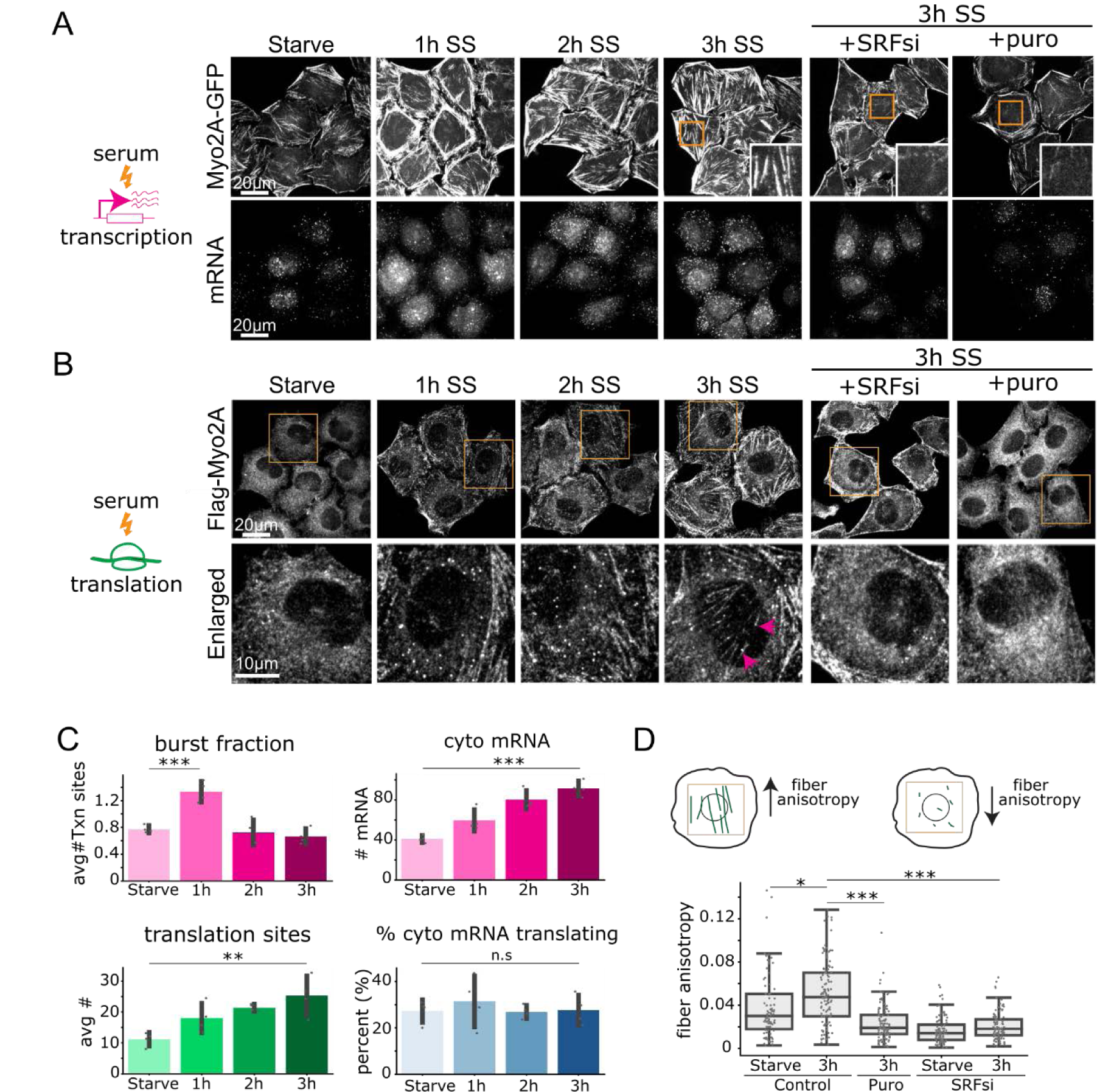
Transcription and translation work together to establish cellular phenotype upon serum stimulation. **A**. Confocal fluorescence images depicting actomyosin cytoskeletal rearrangements and MYH9 mRNAs following serum stimulation (SS). Boxed regions, shown enlarged in insets, exemplify loss of central stress fibers at 3h SS following SRF siRNA or puromycin treatments. **B.** Immunofluorescence images of anti-Flag labeled myosin-2A translation. Boxed regions are shown enlarged in lower panels. Note presence of translation spots associated with a central stress fiber (3h SS, magenta arrowheads). **C.** Quantification of transcription and translation features, values are mean ± SD, n ≥ 156 cells/treatment. **D.** Illustration and measurements of central stress fiber ordering. *p < 0.05, **p ≤ 0.01, ***p ≤ 0.001, n.s, not significant, Welch’s t-test. See also Fig. S4.

Since serum stimulation activates MYH9 transcription, we evaluated whether SRF-dependent transcription and new protein synthesis contributed to actomyosin cytoskeletal reorganization following this stimulation. Transient MYH9 transcriptional activation, and increased levels of its mRNAs and translation sites were all abrogated by silencing of SRF (**Fig. S4C-F**). Notably, while SRF silencing or inhibition of translation with puromycin did not prevent formation of peripheral actomyosin bundles at 1h post serum stimulation (**Fig. S4C**), both treatments disrupted formation of stress fibers over the central portion of cells by 3h post serum stimulation (**Fig. 6A, B**), resulting in reduced linear fiber ordering (**Fig. 6D**). These results suggest that spatial actomyosin stress fiber assembly, within a window of 3h following growth factor stimulus, requires both SRF-dependent transcription and new protein synthesis. Furthermore, the presence of MYH9 translation sites along central stress fibers at 3h post stimulation (**Fig. 6B**, magenta arrowheads) is consistent with a model of localized MYH9 protein synthesis contributing to assembly of these structures. Interestingly, while total levels of MYH9 protein increased marginally following serum stimulation, this remained below statistical significance at the 3h timepoint (**Fig. S4B**). This result is also in accord with a conclusion that new MYH9 protein synthesis may not be required to support global actomyosin activity but rather to localized functions. Changes to myosin-2A protein turnover during this period could contribute to such an outcome.

## Discussion

Homeostatic maintenance of protein levels is an overarching feature of gene expression, but the hidden regulatory mechanisms that maintain homeostasis have been difficult to study and verify. By engineering a novel cell line in which all aspects of MYH9 activity can be visualized in separate colors and with single-molecule precision, we discovered a highly dynamic and multilayered regulatory network lying beneath the apparent stability of myosin-2A expression.

Much like a thermostat maintains temperature, our study provides evidence that MYH9 translation is a key regulatory checkpoint that ensures myosin-2A levels are acceptable. In steady state, nearly every cytoplasmic MYH9 mRNA is translated to meet the demands of the actomyosin cytoskeleton. However, in scenarios when myosin-2A levels become too high, such as when transcription is persistently elevated upon loss of actin depolymerizing proteins like cofilin, translation efficiency is down regulated. Such translational buffering stabilizes protein production to minimize the damaging effects of excessive myosin. Similarly, during cell division, MYH9 translational buffering counters a sharp transcriptional burst. Specific lowering of MYH9 translation during mitosis may safeguard against defects such as impaired karyokinesis, as observed in cofilin-depleted cells with excessive myosin activation ^15^. Conversely, in scenarios that elevate actomyosin assembly, such as the recovery from serum starvation, MYH9 translation aligns with transcription. This is necessary to rapidly remodel the actomyosin cytoskeleton so cellular steady state can be quickly reestablished. Thus, depending on the precise cellular context, our data argue MYH9 translational regulation can either facilitate or buffer the transcriptional response to help ensure myosin-2A levels stay within an acceptable range.

While we were unable to pinpoint the precise molecular mechanism behind MYH9 translational buffering, our study offers some clues. First, the default mode of MYH9 translation seems to be the ON state, since the vast majority of transcripts appear to be translated throughout interphase. Second, the observation that buffering can more than halve the fraction of translating cytoplasmic MYH9 mRNA while leaving their intensities relatively unchanged suggests buffering acts more like a binary on/off switch than a gradual fader. These observation hint at the possible involvement of specific RNA-binding proteins or post-transcriptional modifications that selectively silence MYH9 mRNA. Identifying these specific factors will be an interesting avenue of future research, although a major challenge will be sorting myosin-specific regulators from other targets of the broad SRF/MKL nexus ^29^.

A striking finding of our study is that a just-in-time mode of local protein synthesis appears necessary for assembly of certain actomyosin stress fibers within a 3h period following growth factor stimulation. This result was surprising for at least two reasons. First, growth factor signaling produces rapid stress fiber assembly within 20 minutes of stimulus, suggesting transcription and global new protein synthesis are not required for immediate actomyosin remodeling^26, 27^. Consistent with this view, peripheral actomyosin bundling within 1h of serum stimulus was not lost by perturbations to MYH9 transcription or through protein synthesis inhibition. Secondly, a requirement for new protein synthesis occurred during a period where total MYH9 protein levels did not change significantly. As such, bulk level assays of protein amounts would not have foreshadowed a need for new MYH9 protein synthesis. Nevertheless, loss of central stress fibers at 3h post stimulus, through either inhibition of SRF transcription or protein synthesis, suggested a requirement for both new mRNA transcripts and local translation, similar to what has been observed for actin ^30^. De novo mRNA and protein synthesis of MYH9 and other SRF targets may therefore be a means to specify a distinct population of molecules, without previous modifications, for current localized use.

At the level of transcription, our study sheds light on the elusive molecular pathways that link cofilin inactivation and myosin overactivity. Central to this pathway is MKL, a transcription factor that activates MYH9 via SRF but whose translocation into the nucleus is inhibited by G-actin. According to our data, depletion of the actin depolymerizing proteins cofilin and ADF lowers G-actin levels, leading to MKL nuclear translocation and enhanced transcriptional activation of MYH9. Left unchecked, this leads to excessive myosin contractile forces that, despite the presence of translational buffering, eventually lead to severe morphological aberrations in cells and potentially oncogenic transformations ^31^. Our combined data therefore provide a compelling molecular pathway that would link cofilin-inactivation, higher myosin expression and activity, translational buffering, and disease states.

Having demonstrated the potential of eMyo2AGFP cells, we envision many possible applications. They hold especially great promise for screening purposes, where they can help uncover hidden gene regulatory dynamics. While powerful, there are several caveats to their use worth highlighting. First, it can be technically challenging to detect dimmer myosin-2A translation sites due to the abundance of background signal from mature myosin-2A. While we used the lack of a GFP signal to distinguish translation sites from mature structures, the high background could still be problematic. We therefore recommend measurements of translation efficiency be made in fixed rather than live cells, where FLAG binding sites can be fully saturated and background signal minimized. Second, although we were able to insert mRNA and GFP tags into both MYH9 alleles, we only succeeded in inserting FLAG tags into one allele. This leaves a hidden translation fraction in eMyo2AGFP cells that can only be indirectly assessed. While we believe the hidden fraction behaves similarly to the tagged fraction since we detected translation from nearly half the mRNA, we cannot rule out the possibility that the hidden fraction behaves differently. Third, it remains unclear to what extent our approach can be generalized to other cell lines or other genes. Some cells may be difficult to CRISPR edit and some genes may not tolerate tagging. Keeping these caveats in mind, we believe eMyo2AGFP cells represent a significant advance and should be of broad interest to those wishing to explore the intricacies of gene regulation in the context of cellular perturbations of any kind.

## Methods

### Cell Culture and Drug Treatments

Human HeLa-kyoto cells were obtained from ATCC and maintained in high glucose DMEM supplemented with 10% fetal bovine serum (Atlas). For translation inhibition, cells were treated at 50 μg/ml with puromycin (Sigma) for 20 min or as otherwise indicated. Cells for MKL nuclear-cytoplasmic assessment were treated with 2μM cytochalasin D (Cayman Chemical), 1μM Jasplakinolide (Cayman Chemical) or dimethyl sulfoxide (Sigma) for controls, for 2h prior to fixation for analyses. For serum stimulation experiments cells were starved for 18-24 h in medium containing 0.3% serum and stimulated by replacing with medium containing 15% serum.

### Gene Editing

Sequential rounds of CRISPR/Cas9 gene editing of the endogenous MYH9 gene locus located on chromosome 22 was performed as previously detailed ^32^. Briefly, guide RNAs targeting the start ATG or stop codons of coding exons 1 and 40, respectively, were cloned into plasmid pX330-U6-Chimeric_BB-CBh-hSpCas9. For C-terminal tagging a homology directed repair donor plasmid containing mClover3 (derived from Addgene plasmid 72829), followed by 24xMS2V5 ^33^ was created. For N-terminal tagging a repair donor plasmid consisting of mCherry2 (derived from Addgene plasmid # 72831), flanked by a total of 6X Flag repeat epitope tags was generated. Plasmids for Cas9 and homology repair were transfected to cells with Lipofectamine 2000 (Invitrogen). Successful C-terminal knock-in cells were identified by fluorescence activated cell sorting (FACS) and confirmed by both visual inspection and genomic DNA PCR genotyping, as described ^34^. A clone with two C-terminal knock-in alleles was transfected with Cas9 and N-terminal homology repair plasmids, followed by FACS to isolate dual mCherry2 and mClover positive edited cells. PCR genotyping revealed that all N-terminal knock-in clones had only one of two alleles successfully incorporating 6xFlag-mCherry2. PCR primers to detect the wild type locus could also discriminate a larger product corresponding to the recombinant allele, however, wild type test reactions were optimized and used only to detect the presence of wild type alleles.

### siRNA

Control siRNA oligonucleotides to luciferase and siRNAs cofilin and ADF were as previously described and characterized ^15^. siRNAs to SRF were from Ambion (#13429) and GCGTGAAGA TCA AGATGGA obtained from Qiagen. Cells were transfected 50 nM of siRNAs with Lipofecatmine RNAiMax according to manufacturer’s instructions.

### smiFISH

Cells were fixed for 20 min at room temp in 4% paraformaldehyde (Electron Microscopy Sciences) in PBS, washed in PBS and permeabilized in 70% ethanol overnight at 4 °C. A set of 30 probes spanning the entire coding sequence of human MYH9 were designed using Oligostan software ^35^ and obtained from IDT. Probes were prepared and annealed according to ref 35, mixed with hybridization buffer (Stellaris) and incubated with cells overnight at 37°C. Cells were washed 3x over 30 min in Wash Buffer A (Stellaris), washed once with Wash Buffer B (Stellaris) and mounted for imaging.

### Fluorescence staining

We selected a clone of C-terminally Clover-24xMS2 tagged MYH9 cells that stably expressed Halo-MCP at relatively low levels for visualization of endogenous MYH9 mRNAs. Prior to imaging of mRNAs cells were exposed to 200 nM JF646 HaloTag ligand (Promega) for 20 mins. Cells grown on glass coverslips were fixed in 4% formaldehyde in CBS buffer (10 mM 4-Morpholineethanesulfonic acid, pH 6.1, 138 mM KCl, 3 mM MgCl2, 2 mM ethyleneglycol-bis(β-aminoethyl)-N,N,N′,N′-tetraacetic acid, 0.32 M sucrose) for 20 min at room temperature. mRNAs and cytoskeletal structures were best visualized by inclusion of 0.3% Triton X-100 in the fix buffer. This cytoskeletal fixation procedure enhanced myosin-2A labeling in cytoskeletal structures but resulted in less focal labeling of translation sites as detected by anti-Flag. For best quantitative measures of mRNAs and translation sites, replicate coverslips were fixed with and without inclusion of Triton X-100 to visualize mRNAs and translation sites independently. Translation sites in fixed cells were labeled anti-Flag labeling (FUJIFILM, Wako) followed by Cy3 or Alexa 594 conjugated anti-mouse secondary antibodies (Jackson Immunoresearch and Invitrogen). In live cells, translation sites were labeled by bead loading Flag-Cy3 Fab antibody fragments as previously described ^36^. Cy3-Flag antibodies detect nascent Flag-MYH9 proteins where both mCherry and mClover signals are absent due to the delay in fluorescence protein maturation. Other antibodies were MKL1 (NBP2-45862, Novus Biologicals; sc-32909 Santa Cruz Biotechnology), SRF (66742-1, Proteintech), GAPDH (MAB 374, Millipore), Sun2 (HPA001209, Sigma). DNA was labeled by 4′,6-diamidino-2-phenylindole (DAPI). Mitotic cells were classified based on visual inspection of DAPI labeling.

Cells co-transfected with either control or cofilin/ADF siRNAs and an HA-KDM5B-MS2 translation reporter ^4^ were fixed by the cytoskeletal fixation procedure described above. Cells were labeled for translation by anti-HA antibodies (12CA5, Roche) and with JF646 HaloTag ligand for mRNAs.

### Microscopy

Confocal images were captured on an Olympus IX8 spinning disk microscope with a CSU22 head, equipped with 405, 488, 561 and 640 nm laser lines. Objectives were either 100x/1.40 NA or 60x/1.42 NA. Images were acquired with an iXon Ultra 888 EMCCD camera (Andor) using SlideBook (Intelligent Imaging Innovations). Images of fixed cells were acquired as z-stacks at 0.3 μm intervals.

Cells were plated on glass bottom 35 mm dishes in an enclosed chamber at 37°C and with 5% CO2 for confocal live cell imaging. 11-15 z-planes were acquired at 0.5 μm steps at time intervals between 5 and 10 minutes for evaluations of MYH9 transcription. Live cell translation images were captured as volumes 7-10 z-planes, at 0.65 μm steps and time intervals of 2-10 seconds. Some live cell images were captured on a custom built highly inclined and laminated optical sheet (HILO) microscope ^4^ at similar or faster capture rates.

### Morphometric measurements

Cell, nuclear and cytoplasmic measurements were obtained from 2D segmentations produced using Cellpose software ^37^. Cell masks were obtained using the ‘cyto’ model where image channels marking both cells and DAPI labeled nuclei were utilized. Nuclear masks were generated using the ‘nuclei’ model. Integrated intensity measurements were computed using measures over entire z-stacks from background subtracted images, Measurements of myosin stress fiber ordering through filament anisotropy was done using FibrilTool ^38^. A rectangular ROI covering the central portion, over the entire nucleus, of individual cells was applied to images of ventral actomyosin filaments for measurements.

### Translation site and mRNA Quantification

A subset of maximum intensity z-projections of control or puromycin treated cells, stained with anti-Flag, were used to annotate translation sites for interactive pixel classification machine learning using Ilastik software ^39^. Visually, translation sites were apparent as bright rounded focal spots. All available features from Color/Intensity, Edge and Texture at scales up to smoothing sigma = 5 were selected. Annotations were made for three classes including, background, translation sites and cell. The trained model was used to generate translation site prediction maps for each experimental dataset, that were then segmented in ImageJ to generate centroid coordinates for each detected site. Intensity measurements were made from a central circular ROI with a radius of 4 pixels, using measurements from a 1-2 pixels ROI extension for localized background subtraction. Where indicated, intensity values were normalized between 0 and 1 for all treatments within a replicate.

Images of labeled mRNAs were processed in 3D using FISH-quant v2 ^40^ for detection and quantification of single mRNAs and nuclear mRNA clusters corresponding to transcription sites. A standard FISH-quant pipeline consisted of application of a Laplacian of Gaussian filter to raw mRNA images, followed by spot detection with parameters: voxel size (nm) in 3D = (300, 95, 95), object radius (nm) = (350, 150, 150) and a threshold scaling value of 1.2. Cluster detection values: radius = 350 nm and minimum # spots = 3. Translation sites and mRNAs were mapped to individual cells and nuclei using segmentations of both structures generated by Cellpose. Code from scripts used for image analysis is available at https://github.com/Colorado-State-University-Stasevich-Lab.

### Live cell transcription and translation time series analysis

Transcription sites could be identified as bright pulsatile nuclear spots with intensities greater than three times the mean of single nuclear mRNAs. Visually, cells with transcriptional activity could be readily identified from time series projections of nuclear mRNA distribution. Transcription sites were tracked over time using semi-automated tracking with ImageJ TrackMate ^41^. Frames in which the tracked spot was absent were assigned the position of the last previous observed spot position. Trajectories of spot intensities measured over a 7-pixel radius from spot centroids were used for classification of ON/OFF periods using a Hidden Markov Model (HMM) as introduced previously ^18^. We applied a three-state model in line with an earlier report ^42^. The GaussianHMM function from the python library hmmlearn was used to compute the HMM states for individual transcription site intensity traces.

Tracks of translation site motility were generated with TrackMate and mean squared displacement (MSD) measurements computed using opensource TrackPy.

### Statistical Analysis

Data were tabulated from the results of three to eight experiments, with overlap between datasets for some individual plots such as figures 1E and 5C. P-values were computed using the Welch’s t-test.

## Acknowledgements

We wish to thank Nick Pollock, Ning Zhao and James Bamburg for help with reagents, Luis Aguilera for assistance with computational analyses, Tatsuya Morisaki for scientific discussions and assistance with microscopy and members of the group for their input. This work was supported by grants from the National Institutes of Health (R35GM119728 and R56AI155897) and the National Science Foundation (MCB-1845761).

## Competing interests

The authors declare no competing interests.

## Supplementary Material

**Figure S1.**
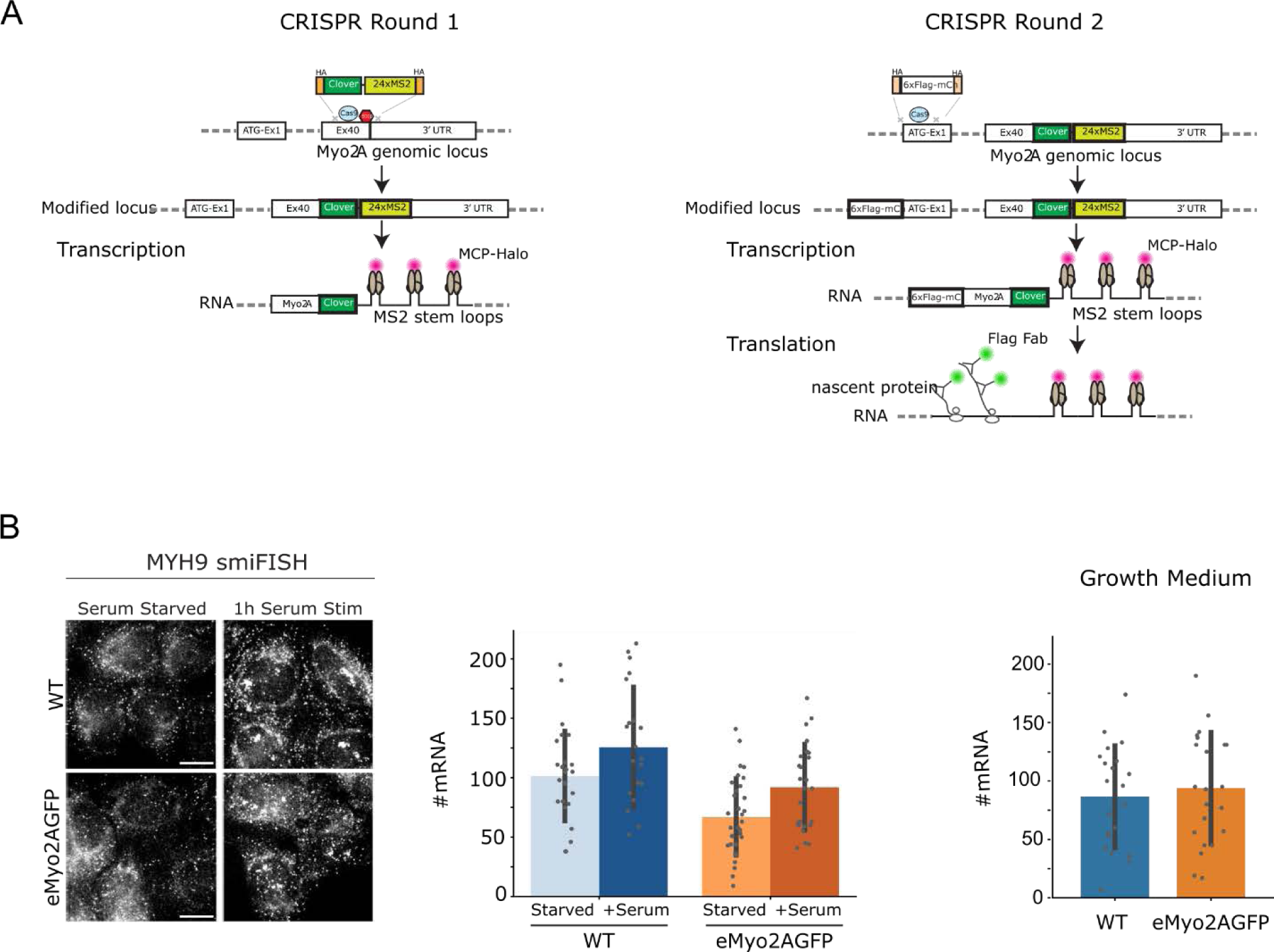
CRISPR strategy to tag endogenous MYH9 and smiFISH analysis of tagged cells. **A**. Illustration of sequential gene editing strategy used to generate eMyo2A GFP cells. **B.** Fluorescence images and quantification of MYH9 mRNA labeled by smiFISH probes that target both tagged and untagged transcripts. mRNAs were assessed under normal growth conditions and following serum stimulation of serum starved cells. Values are mean ± SD. Scale bars 10 μm.

**Figure S2.**
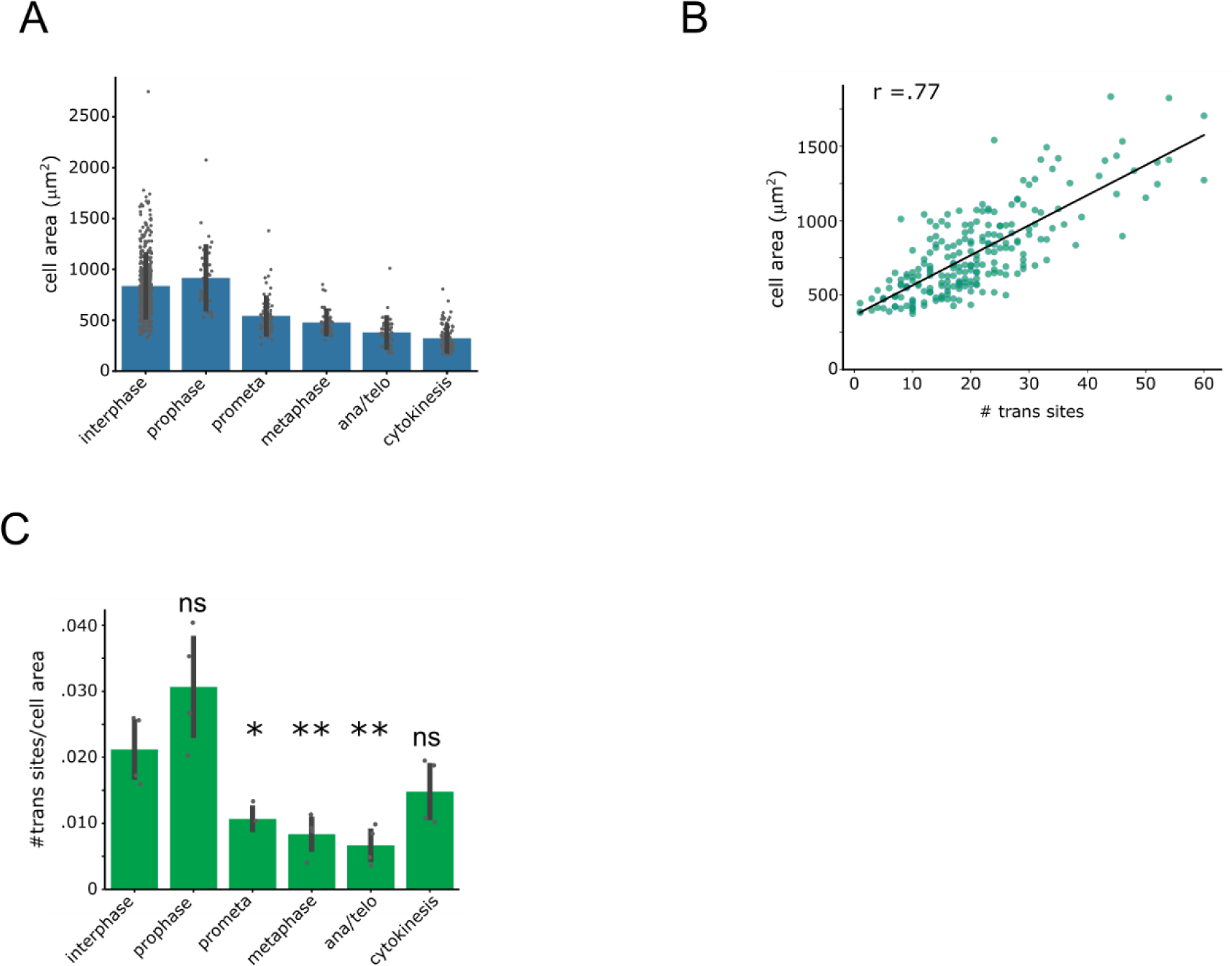
Relationships between MYH9 translation and cell area. **A.** Quantification of eMyo2AGFP surface cell area across the cell cycle, mean ± SD. **B.** Correlation plot of cell area to numbers of MYH9 translation sites, r is Pearson’s correlation coefficient. **C.** Quantification of MYH9 translation sites normalized to cell area across the cell cycle, mean ± SD. *p < 0.05, **p ≤ 0.01, n.s, not significant, Welch’s t-test.

**Figure S3.**
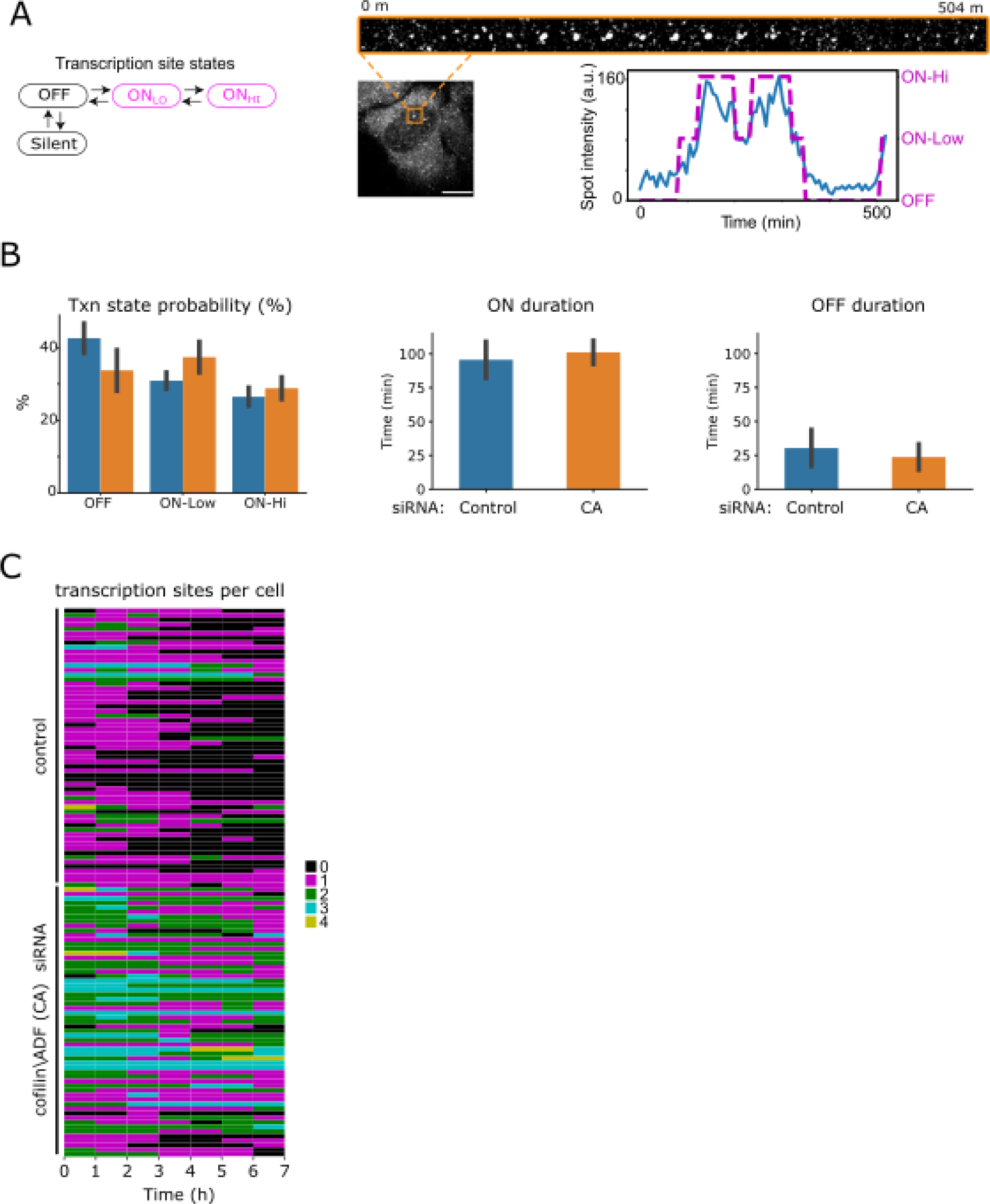
Measurements of MYH9 transcriptional activity in live cells. **A.** Hypothetical transcription activity model and example confocal fluorescence images of Halo-MCP labeled MYH9 transcription foci (boxed region) in a time-lapse series. Scale bar 10 μm. Transcription site intensity trace illustrates classification of on and off states as fit to a hidden Markov model. **B.** Quantification of MYH9 transcription activity dynamics for control or cofilin/ADF siRNA treated eMyo2AGFP cells. Values are mean ± SD. **C.** Heat map of the number of per cell MYH9 transcription sites over time, with each row corresponding to an individual cell.

**Figure S4.**
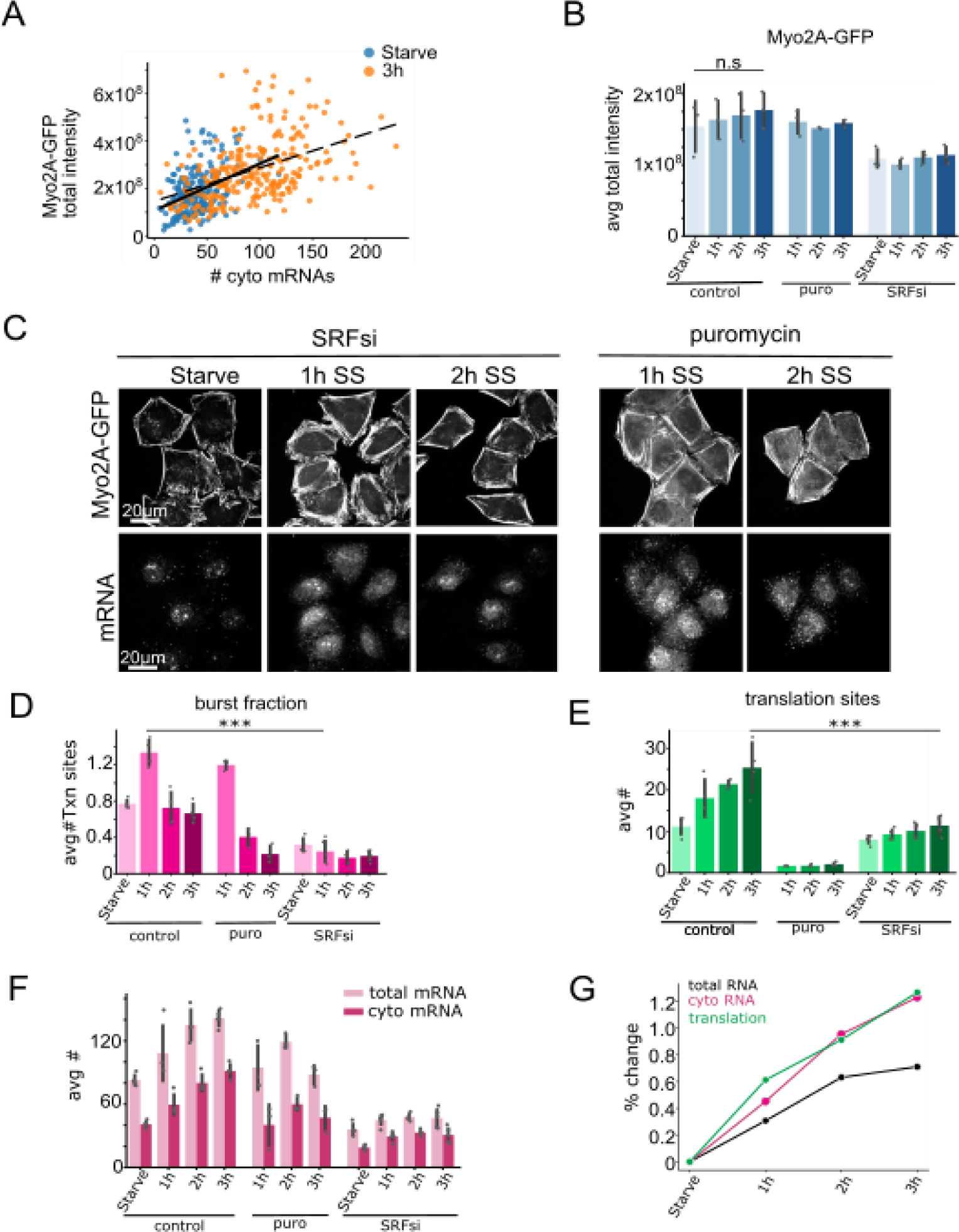
Transcription and translation requirements in MYH9 expression for cytoskeletal response to serum activation. **A.** Correlation plot of integrated Myosin2A-GFP intensity to cytoplasmic MYH9 mRNAs for eMyo2AGFP cells. Solid line shows linear fit to starved condition and dashed line to 3h post serum stimulation. **B.** Quantification of total Myosin2A-GFP intensity. **C.** Confocal fluorescence images depicting actomyosin cytoskeletal rearrangements and MYH9 mRNAs following serum stimulation (SS). **D.** Quantification of the number of per cell MYH9 transcription sites. **E.** Quantification of per cell translation sites. **F.** Quantification of MYH9 mRNAs. **G.** Percentage change plot summarizing data for control treatment from panels E and F. Bar graphs are mean ± SD. ***p ≤ 0.001, n.s, not significant, Welch’s t-test.

## Supplementary Movies

**Movie S1**. Live cell imaging of an eMyo2AGFP cell with myo2A-GFP protein (green) and mRNA (JF646-Halo-MCP, magenta), boxed region illustrates mRNA transcription focus. Images acquired at 1 frame/second. Related to Fig. 1.

**Movie S2.** Live cell imaging of translation from a single mRNA (cyan boxed region, enlarged in spot1) and a mRNA cluster (orange boxed region, enlarged in spot2) in HeLa eMyo2AGFP cells. Translation (trans) and myosin2A protein are labeled by bead loaded Cy3-Flag Fab (green) and mRNAs by JF646-Halo-MCP (magenta). Images are confocal z-stack projections acquired at 0.16 frame (z-volume)/second. Related to Fig. 1.

**Movie S3.** Live cell imaging of a MYH9 transcription activity in an eMyo2AGFP cell with mRNA labeled by JF646-Halo-MCP. Images are confocal z-stack projections acquired at 7 min intervals. Related to Fig. S3

## Notes

### Competing Interest Statement

The authors have declared no competing interest.

### Summary of Updates

Minor text edits.

